# Landmark-Centered Coding in Frontal Cortex Visual Responses

**DOI:** 10.1101/2020.11.04.368308

**Authors:** Adrian Schütz, Vishal Bharmauria, Xiaogang Yan, Hongying Wang, Frank Bremmer, J. Douglas Crawford

## Abstract

Visual landmarks influence spatial cognition [1–3], navigation [4,5] and goal-directed behavior [6–8], but their influence on visual coding in sensorimotor systems is poorly understood [6,9–11]. We hypothesized that visual responses in frontal cortex control gaze areas encode potential targets in an intermediate gaze-centered / landmark-centered reference frame that might depend on specific target-landmark configurations rather than a global mechanism. We tested this hypothesis by recording neural activity in the frontal eye fields (FEF) and supplementary eye fields (SEF) while head-unrestrained macaques engaged in a memory-delay gaze task. Visual response fields (the area of visual space where targets modulate activity) were tested for each neuron in the presence of a background landmark placed at one of four oblique configurations relative to the target stimulus. 102 of 312 FEF and 43 of 256 SEF neurons showed spatially tuned response fields in this task. We then fit these data against a mathematical continuum between a gaze-centered model and a landmark-centered model. When we pooled data across the entire dataset for each neuron, our response field fits did not deviate significantly from the gaze-centered model. However, when we fit response fields separately for each target-landmark configuration, the best fits shifted (mean 37% / 40%) toward landmark-centered coding in FEF / SEF respectively. This confirmed an intermediate gaze / landmark-centered mechanism dependent on local (configuration-dependent) interactions. Overall, these data show that external landmarks influence prefrontal visual responses, likely helping to stabilize gaze goals in the presence of variable eye and head orientations.

**Highlights:**

- Prefrontal visual responses recorded in the presence of visual landmarks
- Response fields showed intermediate gaze / landmark-centered organization
- This influence depended on specific target-landmark configurations

## RESULTS AND DISCUSSION

In this study, we investigated the influence of a static landmark on visual responses in two gaze areas, the FEF and SEF. **Figure 1** shows the visual stimuli that were present before and during the neural responses analyzed in the current study. **Supplementary Figure 1** shows the entire paradigm, including later response periods that we described in previous studies [11–13]. **Figure. 1A** shows an example stimulus configuration where a background landmark (L: a large ‘cross’) first appears, followed by the transient (100 ms) appearance of the target. This cross could appear in one of four spatial (oblique positions) configurations (L1-4) relative to the target stimulus **(Fig. 1B)**. Later, after a delay, monkeys were rewarded for looking at the target stimulus, regardless of any landmark influence **(Supplementary Fig. 1)**. Importantly, animals viewed these stimuli head-unrestrained, to capture the natural complexity of normal gaze behavior. This includes variable torsion of the eyes around the line of sight, which tends to dissociate retina- and world-fixed geometry. This is a challenge for the visual system [14,15], but useful for dissociating these frames experimentally **(Supplementary Fig. 2)**. To account for this, 3D eye orientation was recorded [16], i.e., to precisely calculate the retinal projections of the Target (T) either relative to initial gaze Fixation (TF) or to the Landmark (TL).

**Figure 1:**
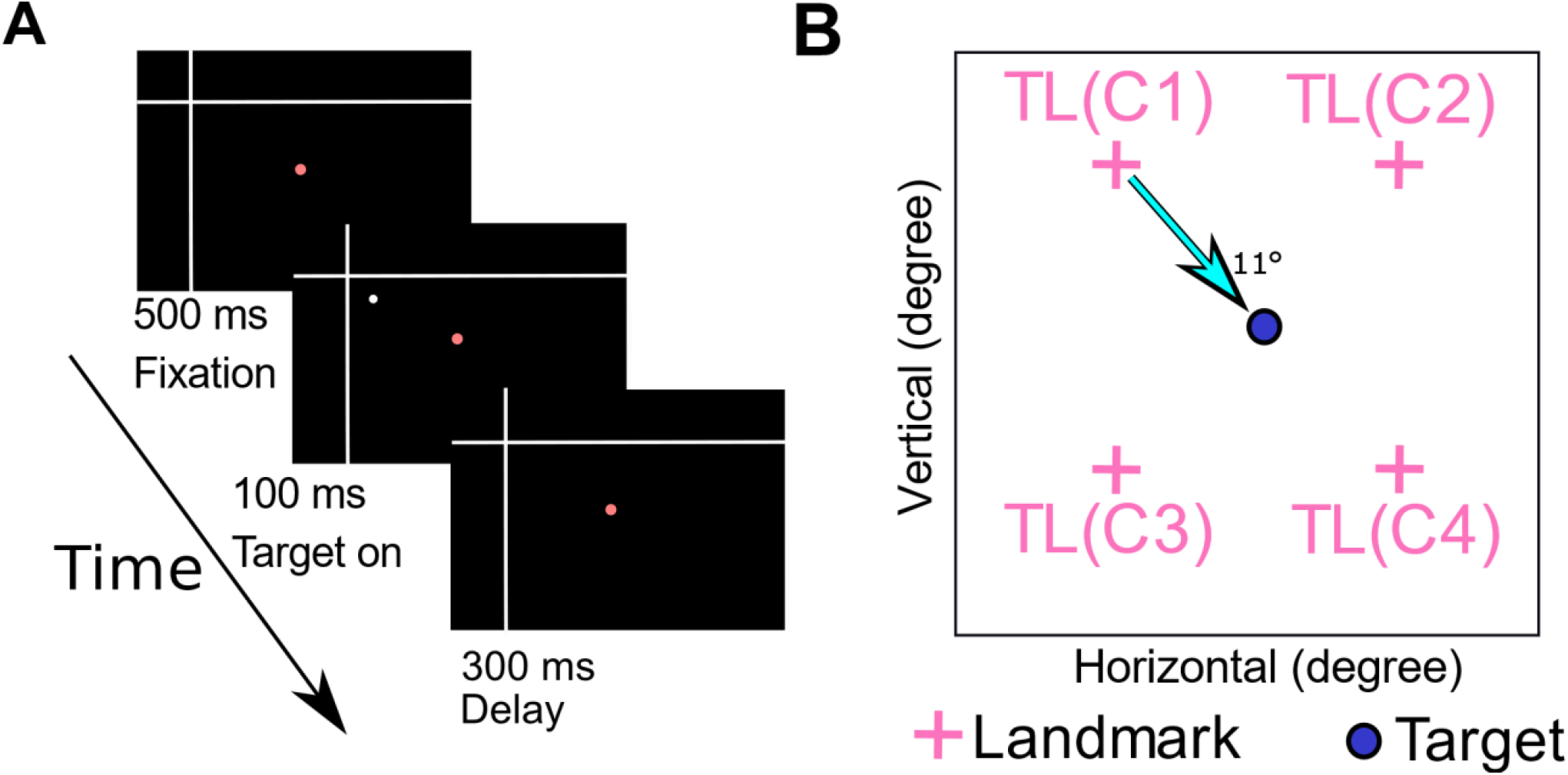
Experimental Paradigm **(A)** Initial stages of the memory-delay / landmark paradigm and its time course. The head-unrestrained monkey starts the trial by fixating a central red dot for 500 ms in the presence of two white intersecting lines (landmark). Then a white dot (target) is flashed (100 ms) in one of four possible locations relative to the landmark, followed by a 300 ms delay. The remaining parts of the paradigm (not analyzed here) are presented in **Supplementary Fig. 1. (B)** Schematic of the four possible target-landmark configurations (TLC1-4).

**Figure 2:**
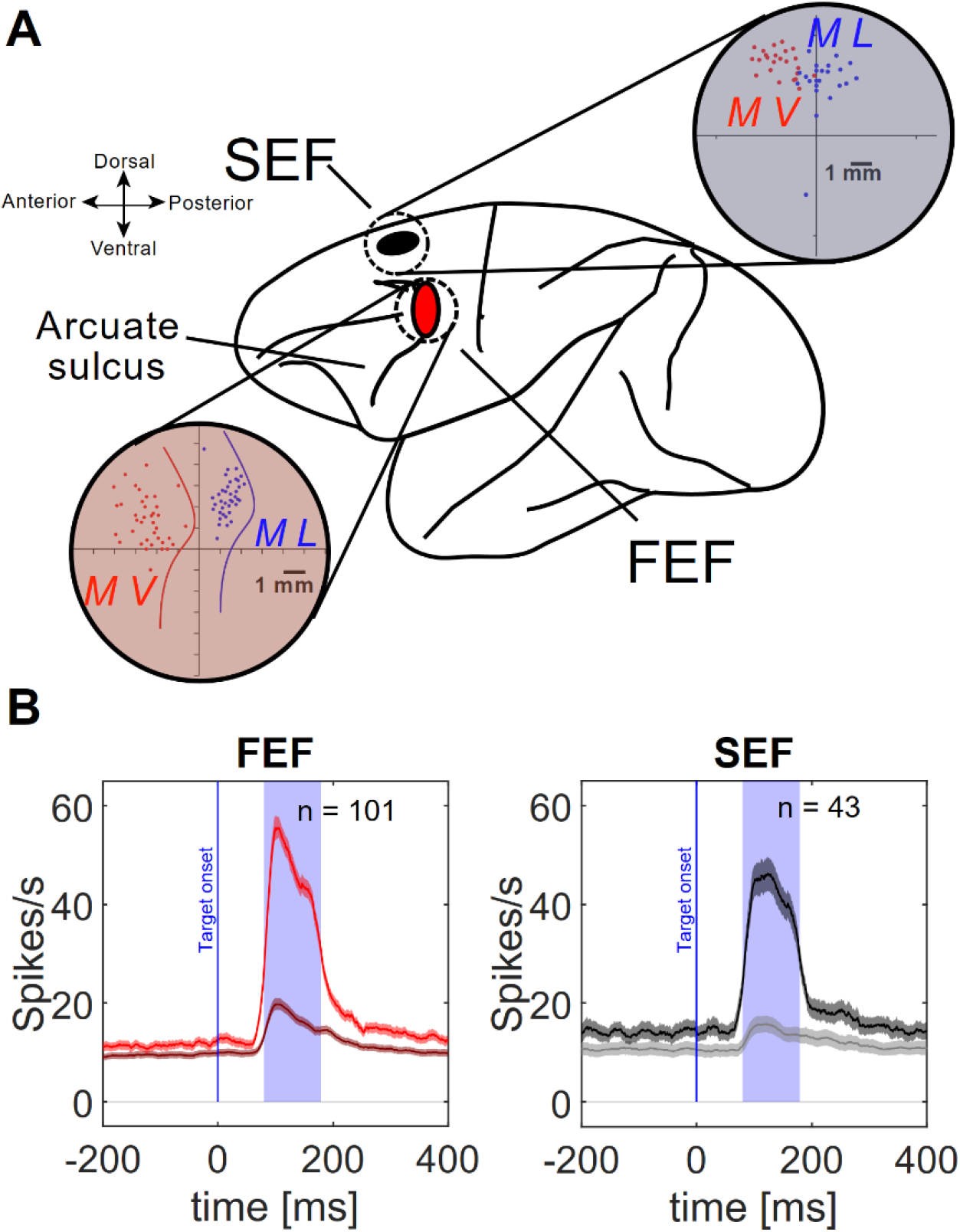
Electrophysiological recordings **(A)** The red ellipse represents the location of the FEF, the black ellipse represents the location of the SEF. The connected red and black disks represent the coordinates of our recording chambers, showing the sites (colored dots) of neural recordings for both monkeys (blue Monkey L, red Monkey V), also confirmed by microstimulation-evoked eye movements. Colored lines in the FEF chamber indicate the location of the arcuate sulcus for both monkeys **(B)** Mean (± SD) of the spike-density plots of the visual responses for all FEF neurons (red) and SEF neurons (black/grey) analyzed in this study. The more robust plots (bright red/black) were derived from the top 10% responses for each neuron (corresponding to the ‘hot’ spot of the neuron’s response field), whereas the dark red / gray plots correspond to data from the entire sample (including the top 10%). The shaded areas show the temporal windows used for sampling data to quantify the visual response (ranging from 80ms to 180ms after target onset).

During neural recordings, targets were presented randomly (one-by-one) throughout each neuron’s response field, while randomly varying the relative landmark configuration, providing a complete dataset for 312 frontal eye field (FEF) and 256 supplementary eye field (SEF) neurons **(Fig. 2A)**. Here, we analyzed the response fields corresponding to the initial visual response to the target, quantified as the number of action potentials within a fixed temporal window after target presentation **(Fig. 2B)**. Off-line, we determined that 102 of the FEF and 43 of the SEF recorded neurons qualified for further analysis by showing statistically significant spatial tuning in their visual response fields (see supplementary methods).

Finally, using our standard methodology [13,17–21] we fit various models against these spatially tuned response fields, focusing on the intermediate reference frames between TF and TL **(Fig. 3A)**. In brief, this involved performing non-parametric fits to the visual response as a function of two-dimensional target location, defined in a specific spatial reference frame **(Fig. 3B)**. The spatial reference frame that yielded the lowest residuals (between the fit and the actual data) was deemed to be the best ‘model’. To test the intermediate reference frame hypothesis, we calculated a series of points along the mathematical continuum between TF and TL **(Fig. 3C1**), yielding response field fits for each of these points [19], where again the fit with lowest residuals ‘wins’ **(Fig. 3C2). Figure 4** provides an example of this analysis for a representative FEF neuron (a similar SEF plot is shown in **Supplementary Fig. 3**).

**Figure 3:**
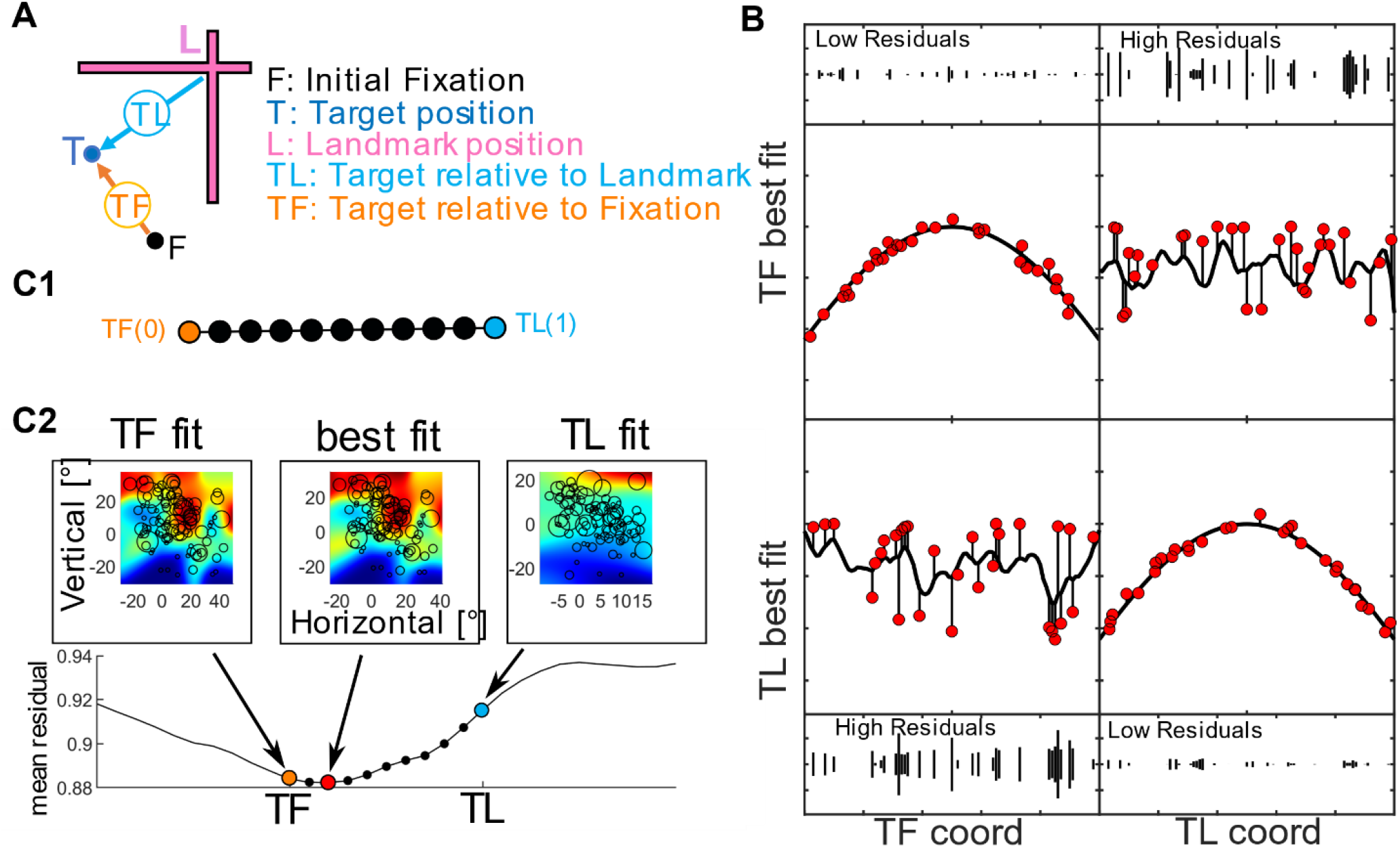
**(A)** Schematic for the spatial parameters of a single trial. The target position (T) is shown in dark blue, the landmark position (L) is shown in magenta, the target position relative to the landmark (TL) is indicated by a cyan arrow, the initial fixation is indicated by the black dot and the target relative to initial gaze fixation (TF) is shown as an orange arrow. All positions were calculated in eye coordinates. **(B**) Schematic of the logic behind the response field analysis. The x-axes represent the spatial coordinate. The y-axes show neural activity. Neural responses from individual trials are represented by the red dots. The black curving lines show the non-parametric fits which do not restrain the response field to a specific (e.g. Gaussian) shape. The upper-left square shows activity from a neuronal response field that is tuned to TF coordinates and plotted relative to TL coordinates, resulting in a good fit with low residuals (difference between the fit and data, shown as vertical lines). The upper-right square shows activity from a neuronal response field that is tuned to TF coordinates, but plotted in TL coordinates, resulting in a poor fit. Conversely, TF coordinates will provide poor fit and TL coordinates will provide a good fit to a neuron tuned to TL coordinates. **(C)** Fitting data to intermediate TL-TF coordinates. **(C1)** Black circles show intermediate steps along the mathematical continuum between TF (represented as 0, orange) and TL (represented at 1, cyan). **(C2)** Upper row: response field fits (heat maps) and responses from individual trials (circles, scaled to neural activity, i.e., number of action potentials in sampling window), plotted in TF, Best Intermediate (0.2), and TL Coordinates. Lower row: mean residuals for fits along the TF-TL continuum, including TF (orange), optimal (red), and TL (cyan) fits.

**Figure 4:**
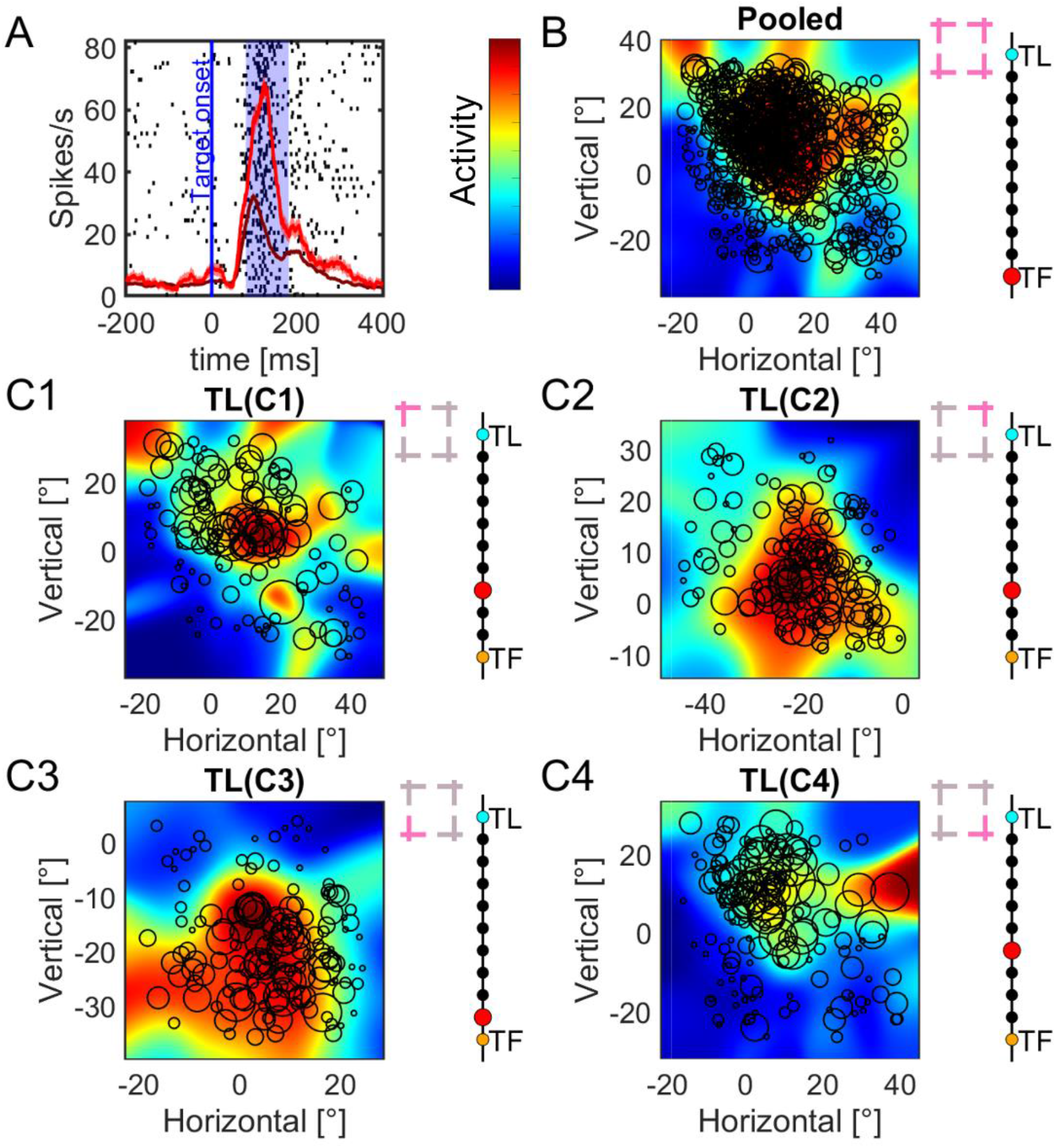
Typical example of a visual response field analysis (in this case for an FEF neuron). **(A)** Raster and spike density plot of the neuron’s activity. The blue line indicates the target onset and the blue shaded area the time window (80ms to 180 ms after target onset) used to quantify the visual response to the target. The dark red line shows the spike density for all trials (including the top 10%) and the bright red line the activity for the 10 % trials with the highest activity in the analysis window. The confidence intervals show the standard error. **(B)** Nonparametric fit of the neuron’s response field after pooling data across all four landmark configurations (indicated by magenta crosses). The color scale on the left indicates the fit to activation ranging from low (blue) to high (red). The black circles indicate responses from individual trials, placed at the location of the target stimulus and the diameter scaled to the response in the sampling window. The pearl thread on the right side indicates the step along the continuum ranging from TF (yellow) to TL (cyan) resulting in the best fit (red). **(C1-C4)** Shows a similar analysis but computed separately for each landmark configuration (TLC1-4, indicated in magenta).

**Figure 4A** represents the raster and spike density plot for the top 10% (red) of neural responses, corresponding to the ‘hot spot’ of the response field [but note that all data (dark red curve) were used for the response field fits]. The blue shaded area (80-180 ms aligned to target onset) corresponds to the temporal sampling window used for the response field plots/fits. **Figure 4B** shows the response field of the neuron, including all trials, plotted against the spatial coordinates derived from the best fit along the TF-TL continuum. Note that here, we pooled across results for all four target-landmark configurations, as indicated by the insets. The black circles superimposed on the colormaps denote the stimulus location and their diameter the visual response for a given trial, whereas the color of the underlying colormap indicates the nonparametric fits to the pooled response field. The best fit for this response field along the TF-TL continuum is denoted by the red dot on the right side of the panel, in this case exactly at TF, seeming to suggest no landmark influence. The results were very similar when we analyzed SEF data in the same way **(Supplementary Fig. 3B)**.

To test the hypothesis that the landmark influence is configuration-dependent we then separated and plotted the response fields for each of the four target-landmark configurations (**Fig. 4C1-4**, where the landmark configuration is highlighted in each of the corresponding insets). Each fit showed a slightly different hot spot (red area), i.e. near the center of the response field in C1 but shifted down or left in the other fits. More importantly, in each case the best fit was shifted toward the TL frame, suggesting an influence of the landmark on the visual response. The same occurred when we separated the SEF data by target-landmark configuration **(Supplementary Fig. 3C1-4)**.

We then performed both types of analyses (pooled and direction-dependent) across the entire populations of spatially tuned neurons in the FEF neurons (N = 102) and the SEF (N = 43). We found no significant difference (Friedman’s ANOVA, FEF: p = 0.59 SEF: p = 0.46) between the distribution of preferred coding along the TF-TL continuum for the four different target-landmark configurations in either the FEF or SEF **(Supplementary Fig. 4)**. Therefore, to obtain one representative fit for each neuron, we recombined the configuration-specific fits by averaging the amounts of TF-TL shift across the four (TLC 1-4) configuration fits for each neuron, thereby standardizing their directional influence.

**Figure 5** contrasts the distributions of the pooled (*left*) and direction-dependent / recombined fits (*right*) for both the FEF (**A**) and SEF (**B**) as ‘violin plots’. For the FEF (**Fig. 5A**) the distribution of fits for the pooled analysis peaked near the TF frame (mean = 0.06 median = 0) with no significant influence of the landmark (p = 1, one sampled sign test), suggesting either that the visual response only codes Target-relative-to-Fixation (TF), or that the landmark influence is obscured when different target-landmark configurations are pooled. Consistent with the latter possibility, the distribution for the configuration-dependent / recombined dataset (right) showed a significant shift toward the TL frame (mean = 0.37, median = 0.4, p = 3.3188*10^−18^ one sampled sign test), suggesting an influence of landmark on the visual responses of FEF neurons. This was also the case for the uncombined target-landmark configurations **(Supplementary Fig. 4A)** and in most individual neurons **(Supplementary Fig. 5A)**. There was also a significant difference between the pooled and the direction-dependent data (two sampled sign test, p = 2.8777*10^−13^), showing that the landmark influence was configuration-dependent.

**Figure 5:**
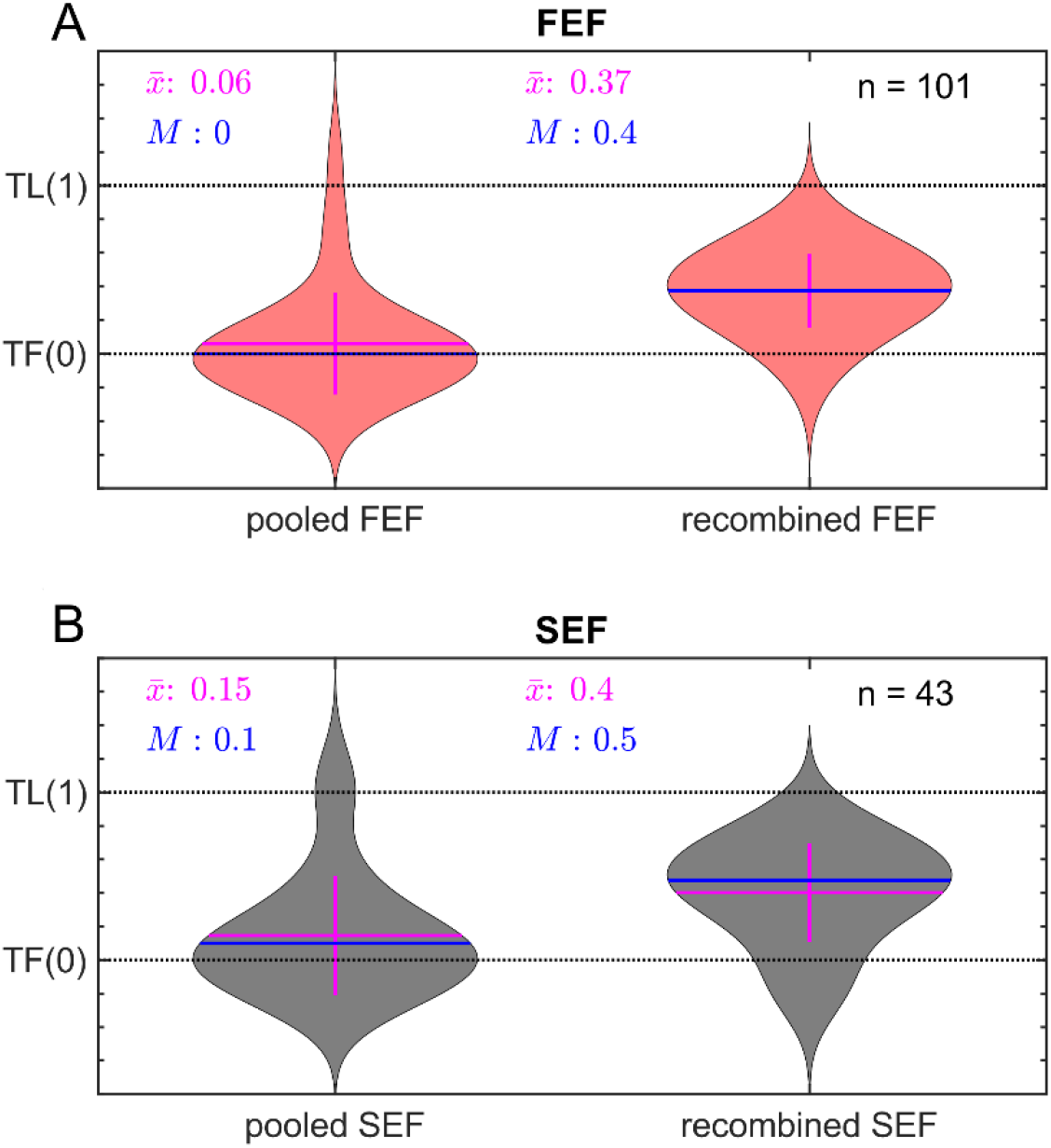
Violin plots of the distributions of best fits along the TF-TL continuum for all spatially tuned FEF neurons (A) and SEF neurons **(B)**. The left side plots show fits made to the pooled data for each neuron; the right-side plots show fits derived separately for each landmark configuration, then recombined by averaging the TF-TL score for each neuron. Blue lines represent the median; magenta lines represent the mean. FEF neurons (A) showed no significant deviation from TF (0) (mean = 0.06, median = 0) in the pooled analysis whereas the separate / recombined distributions (mean = 0.37, median 0.4) were significantly shifted toward TL (1). Likewise, SEF neurons (B) showed no significant shift from TF (mean = 0.15, median = 0.1), whereas the separate / recombined distributions (mean = 0.4, median = 0.5) were significantly shifted toward TL. See text for statistical details.

**Fig. 5B** provides a similar analysis for SEF (n = 43). Overall, we observed the same trends in the SEF as for the FEF. We found no significant shift toward TL (mean = 0.15, median = 0.1, p = 0.13, one sampled sign test) in the pooled analysis (left), i.e., the visual response was best described by the TF model. For the configuration-dependent / recombined analysis (right), a significant shift (mean = 0.40, median 0.5, p = 4.4337*10^−7^, one sampled sign test). Again, this effect was also evident when landmark configurations were tested separately **(Supplementary Fig 4B)**, in most individual neurons **(Supplementary Fig. 5B)**, and there was a significant difference between the pooled data and the direction-dependent data (two sample sign test, p = 1.8217*10^−4^).

In summary, our analysis shows that visual landmarks can influence on the visual code in both the FEF and SEF. More specifically, it caused a shift in the reference frame for coding the target toward the landmark (TL), and further, that this influence was dependent on specific target-landmark configurations. Previously, frontal cortex has been implicated in object-centered coding (one part of an object relative to another) [22,23] and our own studies have shown that prefrontal memory and motor responses are involved in integrating conflicting ego/allocentric cues [11,13]. However, to our knowledge this is the first study that has shown the influence of a stationary, independent, and otherwise behaviorally irrelevant landmark on frontal cortex visual signals to a potential movement target.

It is important to note that this TF-TL shift does not mean response fields shifted toward the landmark. For example, in **Fig. 4C2** the peak of the response field appears to shift *away* from the landmark. Rather, we are talking about a shift in the underlying coordinate frame. In other words, the response of the neuron was dictated by both the location of the target relative to the initial gaze fixation point (TF) *and* the landmark (TL). Based on our response field fitting method, the optimal point was intermediate, i.e., shifted away from TF toward TL. Further, when we pooled data across all TL configurations, their influence was no longer evident, suggesting that the neural influence of the landmark depends on its direction relative to the target (or vice versa: TL). In other words, the visual system seems to implement this influence by forming specific landmark-target linkages rather than a global algorithm that works across configurations. These are likely the reasons why our previous studies failed to find a landmark influence in frontal visual responses [11,13].

The existence of intermediate TF-TL codes in frontal cortex visual responses suggests that the visual system itself is involved in the integration of egocentric (e.g., eye centered) and allocentric (e.g. landmark-centered) cues, but for what purpose? One possibility is that the landmark-centered coding described here helps to compensate for variations of initial eye and head orientation, including eye torsion in space, which is quite large and variable in head-unrestrained conditions **(Supplementary Figure 2A)** and likely helped to distinguish the TF and TL models **(Supplementary Figure 2C)**. It is thought that internal copies of eye and head orientation are used to compensate for these effects [14,15], but visual landmarks can help stabilize noise in these signals [6,12].

Given that the visual system and frontal cortex are reciprocally connected [24–26], the exact site of this influence might be hard to pinpoint. Based on our data, we can speculate that this influence must arise at or before the level of visual inputs to the frontal cortex. The recent finding that some occipital neurons compensate for eye torsion suggests that visual cortex might already contribute to this function [27]. Occipital cortex projects both to parietal cortex, which is associated with egocentric coding, and temporal cortex, which is associated with allocentric coding [9,28–30]. Temporal cortex also projects to parietal cortex [31,32], which in turn projects to both the SEF and FEF [33,34]. Thus, there is ample opportunity for allocentric influence along these pathways.

To our knowledge, there is no precedent for this type of coding in visual responses within the motor system, but what about other neural systems? Place cells and grid cells in the hippocampus account for visual landmarks in the processing of relative location for memory and navigation [35,36]. Recent evidence implies that the hippocampus may derive these signals from as early as the primary and extrastriate visual cortex for coherent encoding of the spatial behavior [37–40]. Altogether these findings and ours suggest that the visual system may play a more sophisticated role in preprocessing allocentric and egocentric cues for other systems, including sensorimotor systems, than previously suspected.

## STAR☆METHODS

Detailed methods are provided in the supplements of this paper and include the following:

- RESOURCE AVAILABILITY
- EXPERIMENTAL MODEL AND SUBJECT DETAILS
- METHOD DETAILS
  ○ Surgical Procedures and recordings of 3D Gaze, Eye and Head
  ○ Behavioral Paradigm
  ○ Behavioral Recordings, Electrophysiological Recordings, Response Field Mapping and Data Inclusion
- QUANTIFICATION AND STATISTICAL ANALYSIS
  ○ Fitting neuronal response fields against Spatial Models
  ○ Pooled vs. Separated analysis
  ○ Intermediate spatial models
  ○ Test for spatial tuning
  ○ Test against randomized background
  ○ Statistical analysis

## RESOURCE AVAILABILITY

**Primary data posted on http://www.yorku.ca/jdc/ (at request / final acceptance)**

## SUPPLEMENTARY METHODS

### EXPERIMENTAL MODEL AND SUBJECT DETAILS

Two female monkeys (*Macaca mulatta*), Monkey V and Monkey L were used for behavioral and neural recordings. All experiments were approved by the York University Animal Care Committee and done in accordance with Canadian Council for Animal Care Guidelines.

### METHOD DETAILS

#### Surgical Procedures and recordings of 3D Gaze, Eye and Head

All experimental procedures were approved by the York University Animal Care Committee and were in accordance with the guidelines of Canadian Council on Animal Care on the use of laboratory animals. The neural data used in this study were collected from two female *Macaca mulatta* monkeys (Monkey V and Monkey L). Both animals were implanted with 2D and 3D search coils. Both search coils had a diameter of 5 mm and were implanted in the sclera of the left eye of the respective animal. The recording chambers for both animals were implanted centered at 25 mm anterior and 19 mm lateral for FEF and 25 mm anterior and 0 mm lateral for SEF. Surgeries were performed as done previously [16]. Underneath each chamber was a craniotomy of 19 mm diameter to allow access to the right FEF and right SEF respectively. During the experiment, the animals were placed in a custom-made primate chair which was modified to allow free head movements. Additionally, the monkey was suited with a vest connected to the primate chair to restrict it from rotating around in the chair. Furthermore, two orthogonal coils were mounted on the head of the monkeys during the experiment. The animal was then placed in the setup which was equipped with three orthogonal magnetic fields. These fields induced a current in each coil. The amount of current induced by one of the fields is proportional to the area of the coil parallel to this field. Thus, allowing to derive the orientation of each coil in relation to the magnetic fields and in turn the orientations, velocities and accelerations of the eye and the head of the animal [16].

#### Behavioral Paradigm

Using a back projector (NEC UM330X), the visual stimuli were presented on a flat screen located 80 cm in front of the animal. The animals were trained on a memory-guided cue-conflict saccade task, where the monkey had to perform a saccade to a remembered target relative to an allocentric landmark (two intersecting lines) that shifted during the memory delay after a mask presentation **(Supplementary Fig. 1)**.

Each trial started with the monkey fixating a red dot located centrally on the screen for 500 ms in the presence of the landmark. Then a white dot serving as visual target was briefly flashed for 100 ms in one of four oblique positions relative to the landmark vertex. Within the context of this paper, each of these target-landmark combinations will be called target landmark configurations [TLC1 (45°), TLC2 (135°), TLC3 (−135°) and TLC4 (−45°)]. For example, TLC1 refers to the Target-landmark configuration where the landmark was present at a 45 ° angle, 11° away from the target. Following a delay of 300 ms a grid-like mask was displayed for 200 ms to occlude visual traces of the landmark. After the offset of the mask, the landmark reappears either shifted (90 %) by 8° in one of eight equally spaced radial directions or not shifted (10%). Following a random delay between 200-600 ms the fixation point disappears acting as a go signal for the animal to initiate a saccade. If the gaze of the monkey landed anywhere in an 8-12° radius around the original target position, the monkey received a droplet of water as reward. This large reward window ensured the monkey is not biased towards either the original target location or the virtually shifted target location fixed to the shifted landmark. Note that all angles mentioned in this section were assumed to be linear. This means an 8° shift in the center of the screen stretches over the same distance on the screen as an 8° shift at the outskirts of the screen.

#### Behavioral Recordings, Electrophysiological Recordings, Response Field Mapping and Data Inclusion

During the experiment, the eye and head orientations in space were recorded at a sampling rate of 1 kHz using the implanted and head mounted search coils respectively. The neuronal activity (Error! Reference source not found.**B)** in the FEF and the SEF was recorded in parallel with tungsten microelectrodes (0.2-2.0 mΩ, FHC Inc.) using the 64 channel Plexon MAP system. To lower the electrodes, the Narishige MO-90 hydraulic micromanipulator was used. The recording sites of the FEF and the SEF were confirmed by using a low-threshold (50 µA) electrical microstimulation while the head was restrained as defined previously [41]. Neurons were mostly searched for while the animal was head-unrestrained scanning its environment. When a reliably spiking neuron was found, the experiment was started. After an initial sampling period for the dimensions of the response field, we presented targets (randomly one-by-one) in a 4 x 4 to 7 x 7 array (each 5-10 ° apart from each other) spanning 30-80° across horizontal and vertical dimensions. We aimed at recording approximately 10 trials/target, so the bigger the response field (and thus the more targets), the more the number of recorded trials were required and vice versa.

For analysis of the visual activity, a fixed 100-ms sampling window was chosen, ranging from 80-180 ms after the onset of the target. Only neurons that showed a significant activation in the sampling window were included in the analysis (Error! Reference source not found.**B)**. Furthermore, trials in which the animals did not successfully fixate on the home position were excluded.

### QUANTIFICATION AND STATISTICAL ANALYSIS

#### Fitting neuronal response fields against Spatial Models

Using the same spatial model-fitting approach that we have employed in this study, we have shown in the SC, the SEF and the FEF that the visual response codes for the target in eye (TF) coordinates [13,17,18]. To examine the influence of the stable visual landmark on the neural activity, here we specifically explored two spatial models **(Fig. 3 A):** 1) the target position in eye coordinates (TF) and 2) the target position in relation to the landmark projected in eye coordinates (TL). The latter is derived by calculating the vector between the landmark and the target and then projecting this vector onto the retina with respect to the 3D-eye position.

To differentiate between different spatial models, they must be spatially separable [17,19]. This variability is ensured by the stimulus design (e.g. random fixation position) and the animal’s natural behavior. Further, opposed to decoding approaches which typically test the set of parameters is implicitly coded in population neuronal activity [42,43], our technique directly tests which underlying spatial model best explains variation in the neuronal activity. The response fields of neurons were fitted against the different spatial models using a non-parametric fit with a Gaussian kernel. To quantify the quality of the fit, the predicted residual error sum of squares (PRESS) statistics was used. These residuals were calculated for each trial by fitting the response field by subtracting the data from the left-out trial and then comparing the activity predicted by the fit for the spatial properties present in the trial and the actual activity measured during the trial. Afterwards these residuals were squared and averaged across each trial to derive PRESS value for a given fit. The bandwidth of the Gaussian kernel was determined for each neuron individually to match the response field’s size, shape, and contour [19]. This was done by calculating the PRESS statistic for each spatial model for all bandwidths between 1 and 15. Then the bandwidth yielding the lowest residuals was deemed as the best fit or spatial model. A schematic of this is displayed in **Figure 3B-C**. Put simply, neural data plotted in the correct reference frame/spatial model would lead to least residuals, e.g. a target-fixed response field would fit best in target-fixed coordinates, whereas in an incorrect frame (e.g. shifted toward Landmark), the data would not fit better, yielding higher residuals. In other words, an intermediate TF-TL point yielded the lowest residuals between the fit and the data, thus best explaining variability in the data. In separate TL configurations, the target-landmark vector was fixed, but variations in initial eye orientation cause this to vary relative to the retina, thus separating TF and TL (**Supplementary Fig. 3)**. This is why our behavioral recordings were important: variations in eye orientation are larger and more variable without head restraint [16], and 3D eye recordings were needed to account for this.

#### Pooled vs. Separate analysis

Previously, when we only analyzed the visual response we did not find any landmark influence in FEF and SEF [11,13], because we pooled the data across all target landmark configurations and did not consider the TF-TL continuum. Here, to quantify the influence of the landmark we contrasted two analysis conditions (pooled and separate). In the pooled condition (also referring to it as direction-independent), all trials of a given neuron were analyzed together, i.e., the response fields were fit with all the data across all the trials for a given neuron. Thus, resulting in a coding preference independent of the TLC **(Fig. 3C)**. However, this can be argued since it can be assumed that the possible influences of the different landmarks are canceled out if all TLCs are combined due to them being oblique. For the separate condition, trials were grouped with respect to the specific TLC **(Fig. 1B)**. i.e., the response fields were fit with the neural data from the trials only corresponding to a landmark in a specific direction (also referred to as direction-dependent analysis). Thus, resulting in four coding preferences/conditions for each neuron (one for each TLC). Since in this pipeline all TLCs are viewed individually, the effects of the landmarks will not cancel out. Note: it is variations in 3D orientation of the eye at fixation that distinguish between the two models **(Supplementary Fig. 3)**.

#### Intermediate spatial models

Our previous results [13,17,18,44] suggested that neuronal response fields do not exactly fit against the canonical spatial models like TF, but rather might best be described by intermediate models between the canonical ones **(Fig. 3C)**. To calculate these intermediate spatial models or continua the linear interpolation between two models was calculated. Further, recently, we have also used this technique to unravel the allocentric influence in gaze activity of the FEF and SEF neurons[11,13]. As mentioned above, here we specifically focused on how the local landmark influence maybe embedded in the activity of visual responses. To this goal, we plotted the sensory response fields along a target-relative to eye to target-fixed to landmark (TF-TL) **(Fig. 3)**. For each continuum there are 31 models along the interpolation axis. Of these 24 models, there are ten frames between the canonical models at 0 and 1; seven beyond model 1; and seven beyond 0.

An example of response field fitting for a continuum between two reference frames (TF and TL) is displayed in **Figure 3C2**. The response fields to the left and right are plotted against the canonical spatial models, i.e., at 0 and 1. The response field at the center corresponds to best spatial frame (shifted 2 steps from TF) for this neuron yielding lowest residuals (red dot).

#### Test for spatial tuning

The method described above assumes the response fields of the neuron are spatially tuned. However, this does not imply that the spatially untuned neurons do not contribute to the code [45–48], but with our technique only the spatially tuned neurons can be explicitly tested. To test for spatial tuning the firing rate data was shuffled over the position data obtained from the best-fitting model [17,19]. The mean PRESS residual distribution (PRESS_random_) of the 100 randomly generated response fields was then statistically compared with the mean PRESS residual (PRESS_best-fit_) distribution of the best-fit model (unshuffled, original data). If the best-fit mean PRESS fell outside of the 95% confidence interval of the distribution of the shuffled mean PRESS, then the neuron’s activity was deemed spatially selective. At the population level, some neurons displayed spatial tuning at certain time-steps and others did not because of low signal/noise ratio. Thus, we removed the time steps where the populational mean spatial coherence (goodness of fit) was statistically indiscriminable from the baseline (before target onset) because there was no task-related information at this time and thus neural activity exhibited no spatial tuning. We defined an index (Coherence Index, CI) for spatial tuning. CI for a single neuron, which was calculated as [49]:

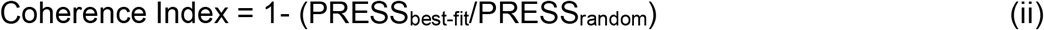

If the PRESS_best-fit_ was similar to PRESS_random_ then the CI would be roughly 0, whereas if the best-fit model is a perfect fit (i.e., PRESS_best-fit_ = 0), then the CI would be 1. We only included those neurons in our analysis that showed significant spatial tuning.

#### Test against randomized background

To verify that the changes in the coding preference in the separated analysis are correlated with the TLC we tested each individual neuron’s coding for each of the four TLC groupings against a background. The background was derived by shuffling the TLC groups and then calculating the resulting coding preference. This ensures that possible effects on the coding preference by the specific grouping is removed allowing for a quantification of the influence of the grouping.

##### Statistical analysis

All statistical analyses were performed using MATLAB R2019b. We assumed a significance level of p < 0.05.

#### KEY RESOURCE TABLE

| REAGENT or RESOURCE | SOURCE | IDENTIFIER |
| --- | --- | --- |
| **Deposited Data** |  |  |
| Raw and analyzed data | This paper |  |
| **Experimental Models: Organisms/Strains** |  |  |
| Macaca Mulatta |  | N/A |
| **Software and Algorithms** |  |  |
| Plexon Offline Sorter | Plexon |  |
| Analysis Scripts | This paper |  |
| Matlab R2019b | Mathworks |  |
| **Other** |  |  |
| Narishige MO-90 hydraulic micromanipulator | Narishige |  |
| Tungsten microelectrodes (0.2-2.0 mΩ) | FHC Inc. |  |
| NEC UM330X | NEC Display Solutions |  |
| 64 channel Plexon MAP system | Plexon |  |
| Lab made coil system | Crawford et al. 1999 | N/A |

## Conflict of Interest

The authors declare no conflicts of interest

## Acknowledgement

This project was supported by a Canadian Institutes for Health Research (CIHR) Grant and the Vision: Science to Applications (VISTA) Program, which is supported in part by the Canada first Research Excellence Fund, and by Deutsche Forschungsgemeinschaft (DFG: IRTG-1901, RU-1847, and CRC/TRR-135, project number 222641018). VB, XY, and HW were supported by CIHR and VISTA. AS was funded by DFG. JDC is supported by the Canada Research Chair Program.

## Supplementary Figures and Analysis

**Figure 1:**
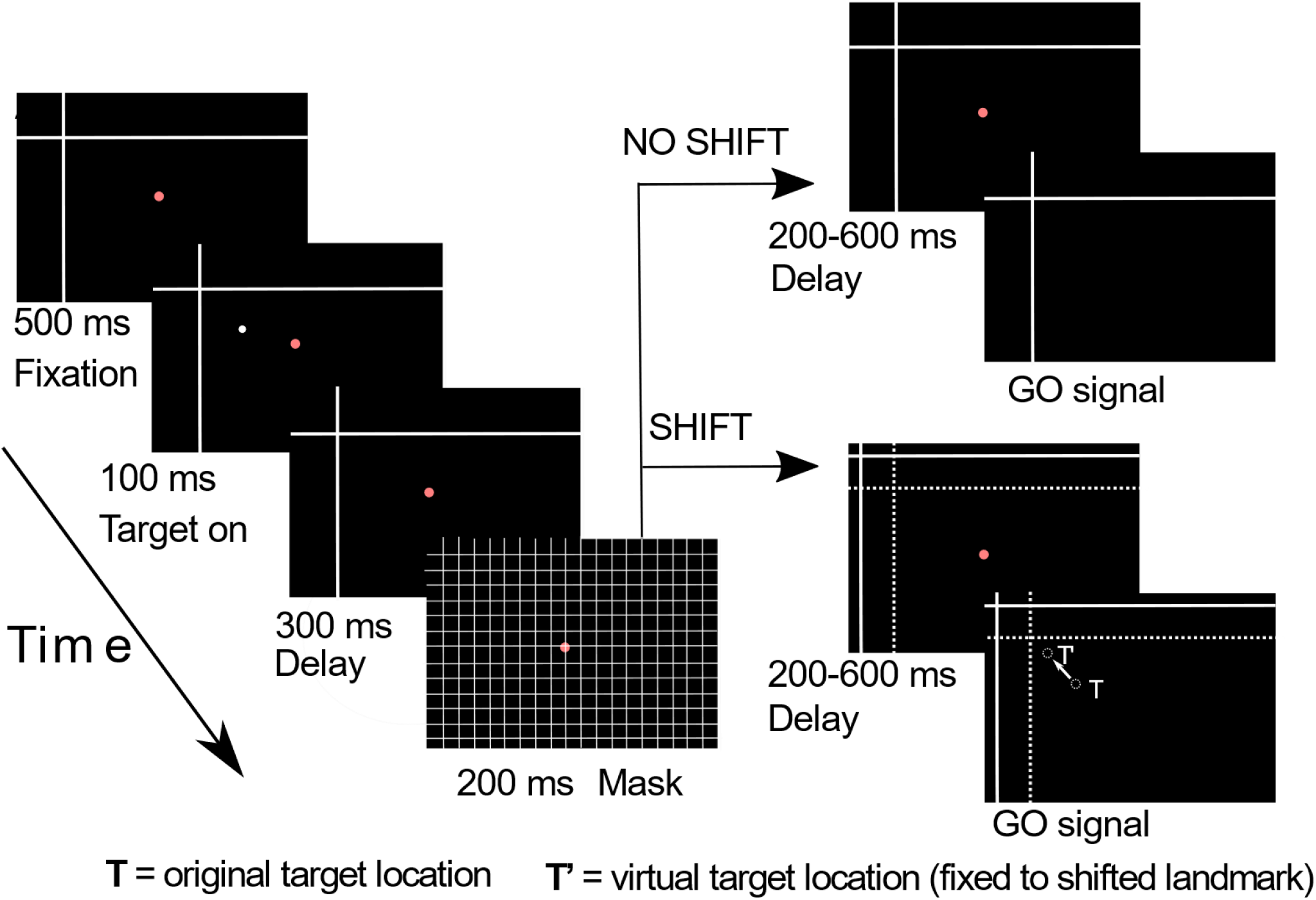
(A) Complete memory-delay / landmark paradigm, showing cue-conflict and saccade ‘go’ signal that occurred later in the time course (not analyzed here). The monkey starts the trial by fixating a central red dot for 500 ms in the presence of two white intersecting lines (landmark). Then a white dot (target) is flashed (100 ms) in one of four possible locations relative to the landmark, followed by 300 ms delay and a grid-like mask (200 ms). After mask offset the landmark either shifted in one of eight radial locations from the original location (or not shifted). Then after an additional delay period (200 – 600 ms) the fixation point was extinguished which served as a go signal for the animal to saccade to the remembered target location. The animal was rewarded for landing its gaze (G) within a circle of radius 8-12° around the original target. Gray box: Portion of the stimulus associated with the visual response.

**Figure 2:**
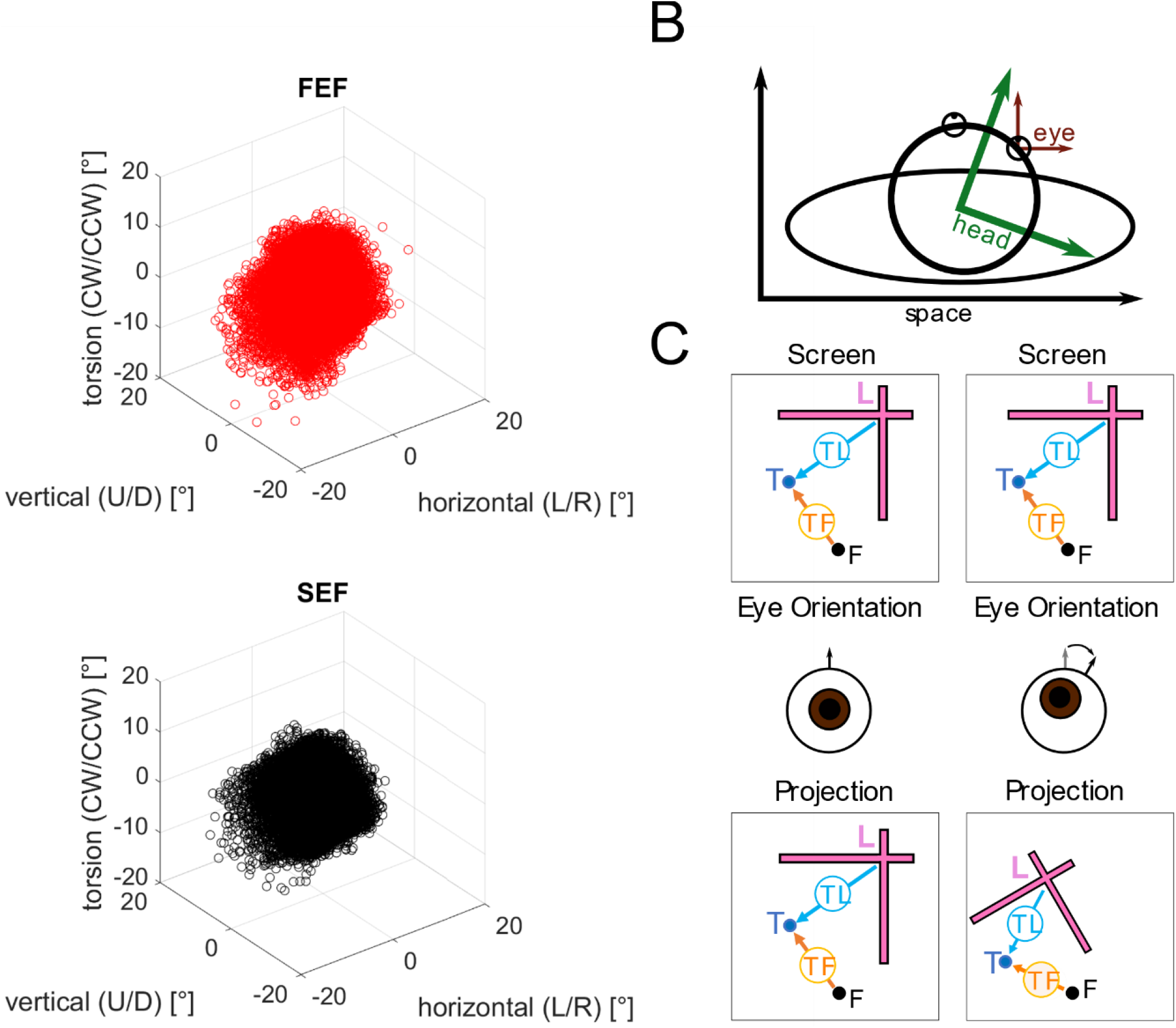
**(A)** Scatter plot of eye orientation at initial fixation position. Orientation is given by the composite of eye-in-head and head-in-space orientation. Every trial used in the analysis is shown. **(B)** Schematic of how different rotation in different coordinate systems interact with each other. As one can see a change in head-in-space orientation induces changes for the eye-in-space orientation. **(C)** Schematic of how eye orientation in space influence projection of the stimulus. The top row represents the screen, the middle row show a schematic eye orientation and the bottom row show the resulting projection based on this eye orientation. As one can see a tilted and rotated eye result in a rotated and compressed projection of the presented scene. Thus, leading to different projected TL vectors for the same TLC. The amount of compression is greatly exagarated for demonstrative purposes.

**Figure 3:**
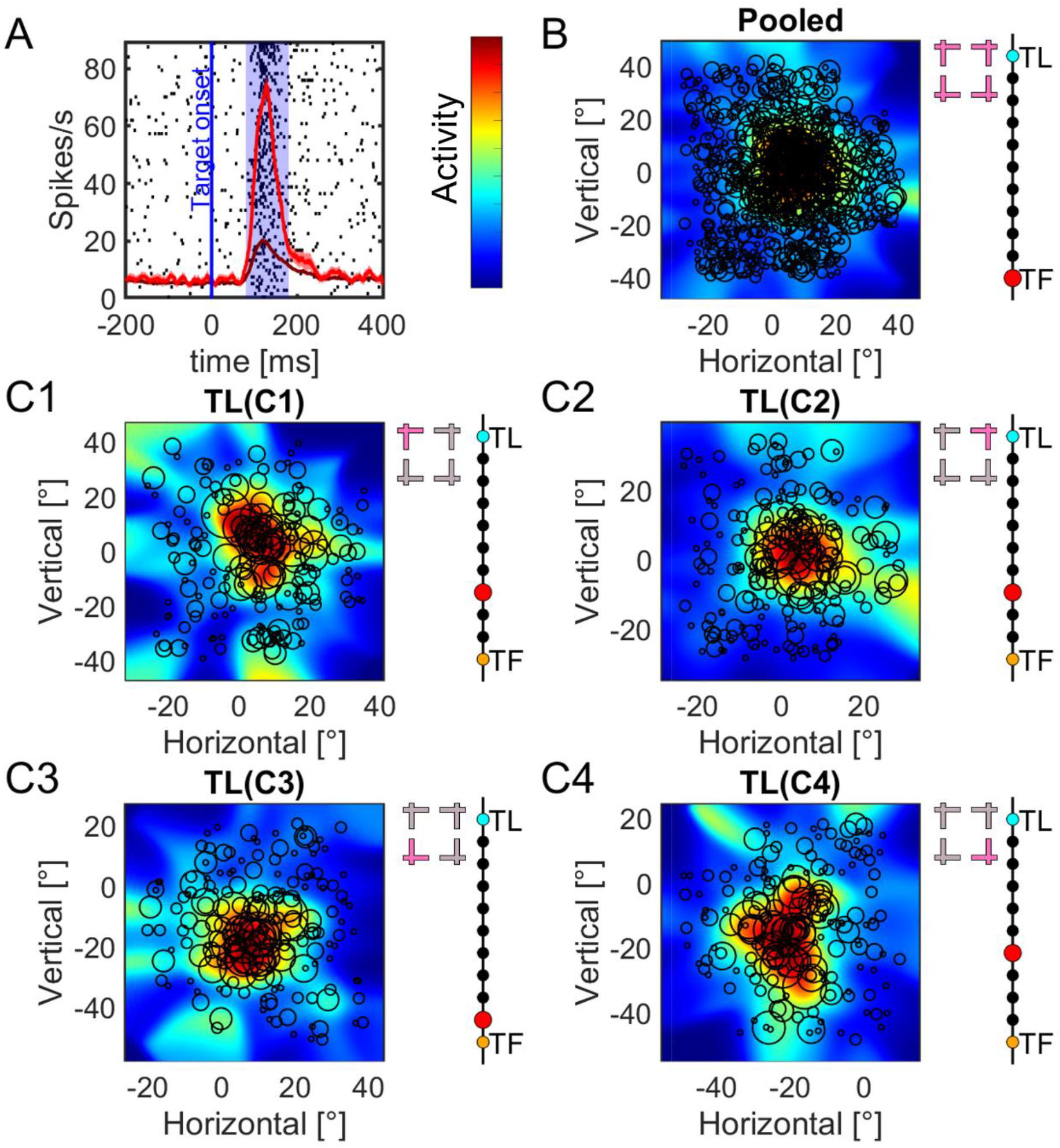
presents a typical example of a visual response analysis (for SEF neuron). **(A)** Shows the raster of the neuron activity. The blue line indicates the target onset and the blue shaded area the time window used for further analysis. The dim red line shows the activity density for all trials, the red line the activity for the 10 % trials with the highest activity in the time window between 80 ms to 180 ms after target onset. **(B)** Nonparametric fit of the neurons response field. The color scale on the left indicates activation ranging from low (blue) to high (red). The black circles position show the position of a trial in the feature space the size of the circle the corresponding neuronal activation in this trial in the analysed time window. The pearl threat on the left-hand side indicates the optimal step (red) along the continuum ranging from TF (yellow) to TL (cyan). **(C)** Shows the same analysis with the same figures convention as (B) but here the trials are splitted according to their respective target-landmark-configuration (TL(Cn)).

**Figure 4:**
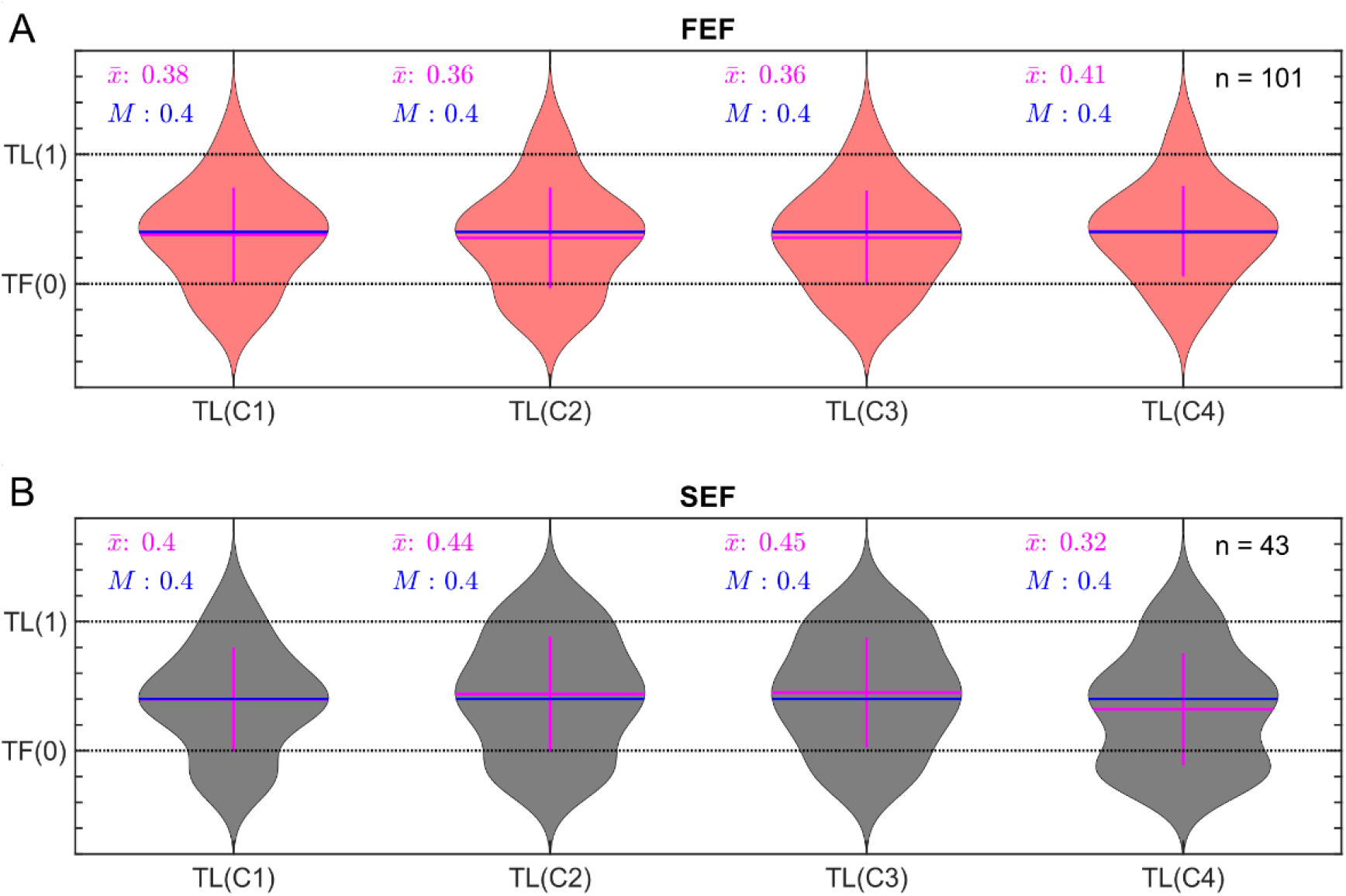
Violin plot of coding preference of neural population along the TF-TL continuum. The top row/red plots show FEF neurons. The lower row/black plots show SEF neurons. Blue lines represent the median magenta lines represent the mean. **(A)** Best fit distribution for FEF neurons along the TF–TL continuum for each Target-Landmark configuration TLC1-4 (mean = 0.38, 0.36, 0.36, 0.41, median = 0.4, 0.4, 0.4, 0.4 respectively). **(B)** Best fit distribution for SEF neurons along the TF–TL continuum for for TLC1-4 (mean = 0.40, 0.44, 0.45, 0.32, median = 0.4, 0.4, 0.4, 0.4, respectively).

**Figure 5:**
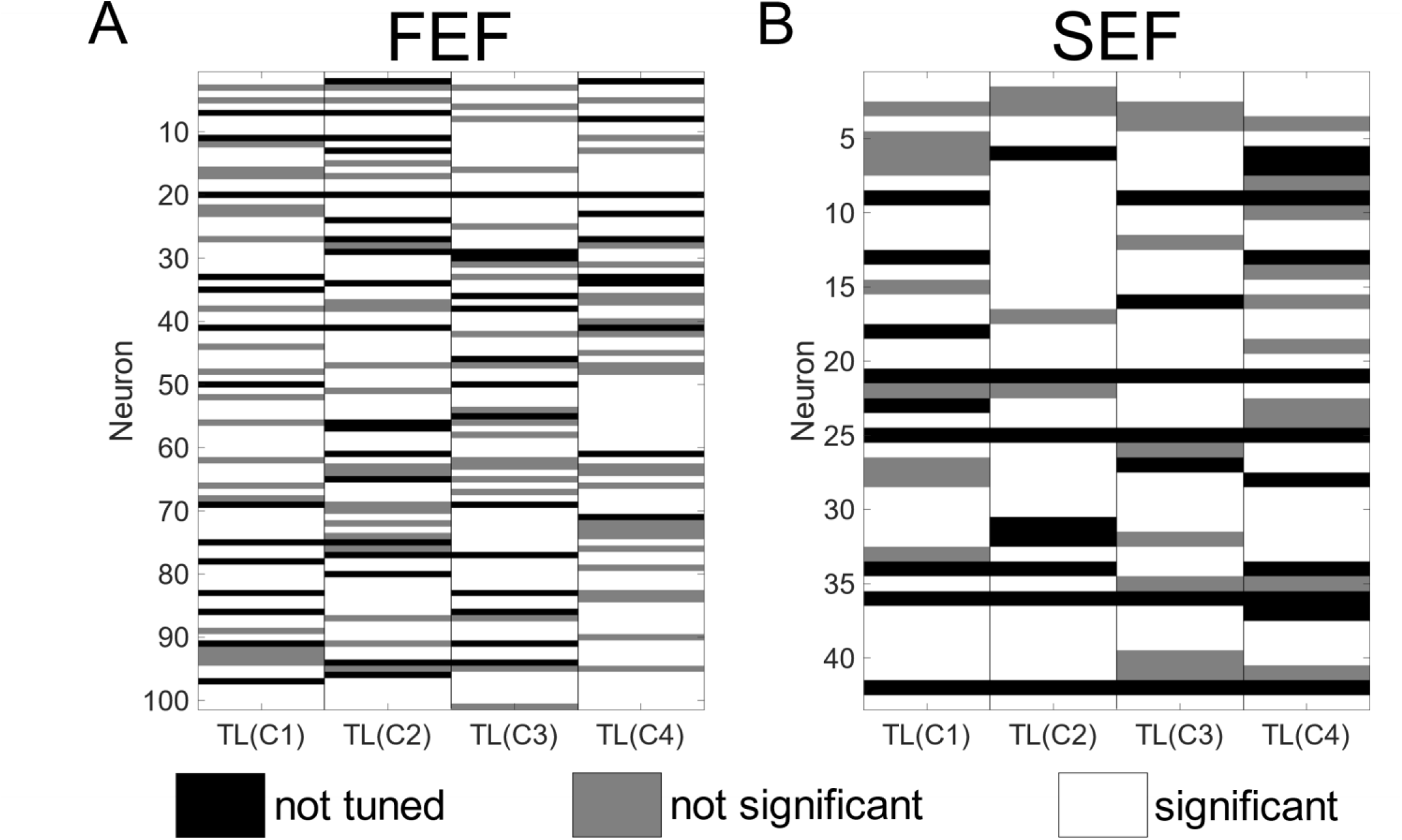
Neuron-by-neuron significance test for TF-TL shift, broken down by Landmark configuration (TLC 1-4). Left: FEF neurons Right: SEF neurons. The y axis identifies the neuron, the x axis the represents the TLC. Black cells are significantly different. From their respective background describes in methods. Red cells are not significantly different. White cells are untuned neurons.

## Notes

### Competing Interest Statement

The authors have declared no competing interest.

